# Copper induces p53 unfolding in cancer cells, resulting in invasion and chemoresistance

**DOI:** 10.1101/2024.11.07.622423

**Authors:** Yannick von Grabowiecki, Steven J Bell, Hannah L Richards, Ellie J Massey, Zoe Newson, Emma Tarrant, Matthew S Morrison, Melanie Galvin, Kathryn L. Simpson, John Le Quesne, Lydia Aschauer, Patricia AJ Muller

## Abstract

Copper has been shown to play an important role in cancers, enhancing cell proliferation, remodelling the microenvironment and enhancing the function of oncogenes. Here we show that copper turns the tumour suppressor p53 protein into a pro-oncogenic protein.

Mutations in the *TP53* gene often lead to expression of various mutant p53 proteins. Mutant proteins often lose some or all wild type function and gain novel pro-tumourigenic functions, which can be partially attributed to protein unfolding. Here we show that copper accumulates in tumours, unfolds WTp53 and promotes chemoresistance. Unfolding of WT p53 results in interaction with mutant p53-specific interacting proteins TAp63 and Ago2 and promotes invasion. Interestingly, partially-functional p53 mutants that are frequently observed in lung cancers, are more affected by copper than WTp53. Our results suggest that small increases in copper impair WT p53 function and augment mutant p53 function, which could partially be restored by the chelator Trien.

## Introduction

The p53 tumour suppressor plays a central role in the cell in processes such as cell cycle arrest, DNA repair or apoptosis preventing the build-up of DNA damage in cells (Bieging et al., 2014). The importance of this protein is underpinned by the fact that somatic mutations of the TP53 gene occur in more than 50% of human cancers (Olivier et al., 2010; Stracquadanio et al., 2016; Wang and Sun, 2017). The majority of TP53 mutations are located in the DNA-binding domain and represent missense mutations, leading to mutant p53 expression(Olivier et al., 2010). In general, mutant p53 proteins can be classified in two groups: DNA-contact mutations or conformational mutations, depending on whether the mutation affects residues involved in DNA binding or if the tertiary structure is affected. (Bullock et al., 2000; Joerger et al., 2005).

Mutations in the p53 gene (*TP53)* lead to loss or reduced function of the protein (Hachiya et al., 1994; Willis et al., 2004). In addition, mutant p53 can acquire new oncogenic functions, gain-of-functions (GOFs), which include increased invasion and metastasis, increased cell proliferation and increased chemo-resistance(Muller and Vousden, 2014). These GOFs are promoted by binding of mutant p53 to other proteins (Aschauer and Muller, 2016; Muller and Vousden, 2014). Well-described interaction partners of mutant p53 are its family members p63 and p73, whose tumour suppressive functions are inhibited by this interaction (Di Como et al., 1999b; Gaiddon et al., 2001; Irwin et al., 2003; Muller et al., 2009; Strano et al., 2002; Xu et al., 2011). Other interaction partners are microRNA machine proteins, such as Ago2 (Krell et al., 2016).

These novel interactions of mutant p53 are often a result of structural distortion and reduced flexibility of the protein found in both contact and structural p53 mutants(Olotu and Soliman, 2017). Noteworthy, the Wild Type p53 (WT p53) protein is considered unstable as 50% of the protein is unfolded in the absence of stabilising partners at 37°C (Bell et al., 2002; Cañadillas et al., 2006). In addition, hypoxia unfolds p53 (Gogna et al., 2012) and correct folding requires the presence of the chaperonin containing tailless complex polypeptide 1 (CCT) complex (Trinidad et al., 2013). Structural studies of p53 revealed that the interaction with a zinc ion has an essential role in stabilising the DNA core domain of p53 (Joerger et al., 2005). Thus, chelation of zinc leads to unfolding of WTp53 and reduced DNA-binding activity(Hainaut and Milner, 1993b; Verhaegh et al., 1998). Divalent metals such as copper, were shown to unfold WTp53 (Hainaut and Milner, 1993b; Meplan et al., 1999a; Meplan et al., 1999b; Tassabehji et al., 2005). Copper can affect p53 function in different manners: cause DNA damage directly or indirectly via induction of reactive oxygen species (ROS), alter cellular localisation or expression of p53, or unfold the p53 protein (Phatak and Muller, 2015a). The extent to which copper contributes to changes in p53 function in cancer cells remains elusive.

Many studies have demonstrated an important role for copper in cancer cell metabolism and promoting oncogenic functions. Most prominently, the high energy demand for cancer cells requires copper to fuel metabolism as copper is a co-factor for enzymes such as cytochrome C (Ruiz et al., 2021). Additionally, copper promotes invasion and metastasis, which could in part be the result of its role in epithelial to mesenchymal transition (EMT) (Focaccio et al., 2024). Copper levels in serum, urine or blood were reported to be higher in patients with diverse malignancies compared to normal individuals (Gupte and Mumper, 2009). In addition, biomarkers such as expression of the copper importer *SLC31A1* or the copper-responsive genes metallothionein genes have been used to associate high copper levels to cancers, cancer progression, metastasis or treatment response (Kong et al., 2023; Qi et al., 2023; Werynska et al., 2013; Yun et al., 2022). However, only a handful of reports have examined actual copper levels within cancer cells, compared to adjacent tissue or serum levels or compared to normal tissues (Díez et al., 1989; Shanbhag et al., 2021).

Occupational exposure (metal working, painting or stonemasonry) and tobacco are considered the highest risk for developing small cell lung cancer (Soleimani et al., 2022). The fumes that one is exposed to in these professions as well as the smoke from cigarettes are known to contain high copper levels. Associations between lung cancer and increased copper levels exist, but whether copper is accumulated in the cancer cells, whether p53 levels are impaired and how copper promotes lung cancer cell formation are still not fully understood.

In this study, we show copper accumulation in tumour cells of small cell lung cancer patients. We investigated the folding status of WTp53 in response to copper in several cell lines and we show that copper-unfolded WTp53 interacts with the mutant p53 binding proteins p63, p73 and Ago2. Furthermore, we reveal that copper exposure induces invasion in a p53-dependent manner and causes chemoresistance. Interestingly, lung cancer-specific partially-functional mutants are more vulnerable to copper unfolding than WTp53. Our data suggest that copper-dependent unfolded p53 not only becomes inactive, but acquires a GOF that drives tumour formation and causes chemoresistance.

## Results

### Copper unfolds p53 and causes loss of p53 transcriptional function

To understand the relationship between p53 function and conformational changes, we noted that copper (CuSO_4_) is very potent in unfolding p53 (Figure 1A). The folding status of p53 can be assessed by two conformation-specific p53 antibodies. The PAb1620 (1620) antibody only recognizes the wild type conformation (Wang et al., 2001) and the PAb240 (240) only recognises only a “mutant”-like/denatured conformation of p53 (Stephen and Lane, 1992). Using these antibodies in immunoprecipitation on native lysates of HEK293T cells, we observed a dose-dependent increase in PAb240 binding and a decrease in PAb1620 binding (Figure 1A), clearly showing unfolding of p53 from as low as 5 µM of CuSO_4_. Using in-cell westerns and a fixation technique that retains proteins folding, we validated p53 unfolding in response to copper (Figure 1B, Figure S1A). These data confirm that we are looking at intracellularly unfolded p53, rather than an artefact of copper being released from other proteins upon lysis. HEK293T cells contain the SV40 large tumour antigen, which binds at least partially inactivates WT p53 (Sheppard et al., 1999). To study p53 function in cells with a fully functional p53 response, we used the bronchial epithelial lung cell line Beas-2B and the non-small cell lung cancer cell line A549. Similar to HEK293T cells, we could detect a dose-dependent decrease in p53 folding upon copper incubation (Figure 1C and 1G). Although some literature speculates on loss of p53 function in response to unfolding, remarkably few studies actually show this (Phatak and Muller, 2015b; Tassabehji et al., 2005). Using the PG13 luc (WTp53 binding sites) luciferase construct that contains p53 binding elements upstream of the luciferase coding sequence, we could observe a decrease in p53 transcriptional function in Beas-2B and A549 cells upon copper exposure (Figure 1D and H). Increased intracellular copper levels were confirmed by ICP-MS that measures total copper levels (Figure 1E and I), as well as a MRE luciferase reporter assay in which metal response elements are positioned upstream of the luciferase coding sequence and therefore reflects intracellular copper signalling (Figure 1F and J). These data indicate that p53 activity is reduced upon intracellular copper accumulations.

**Figure 1.**
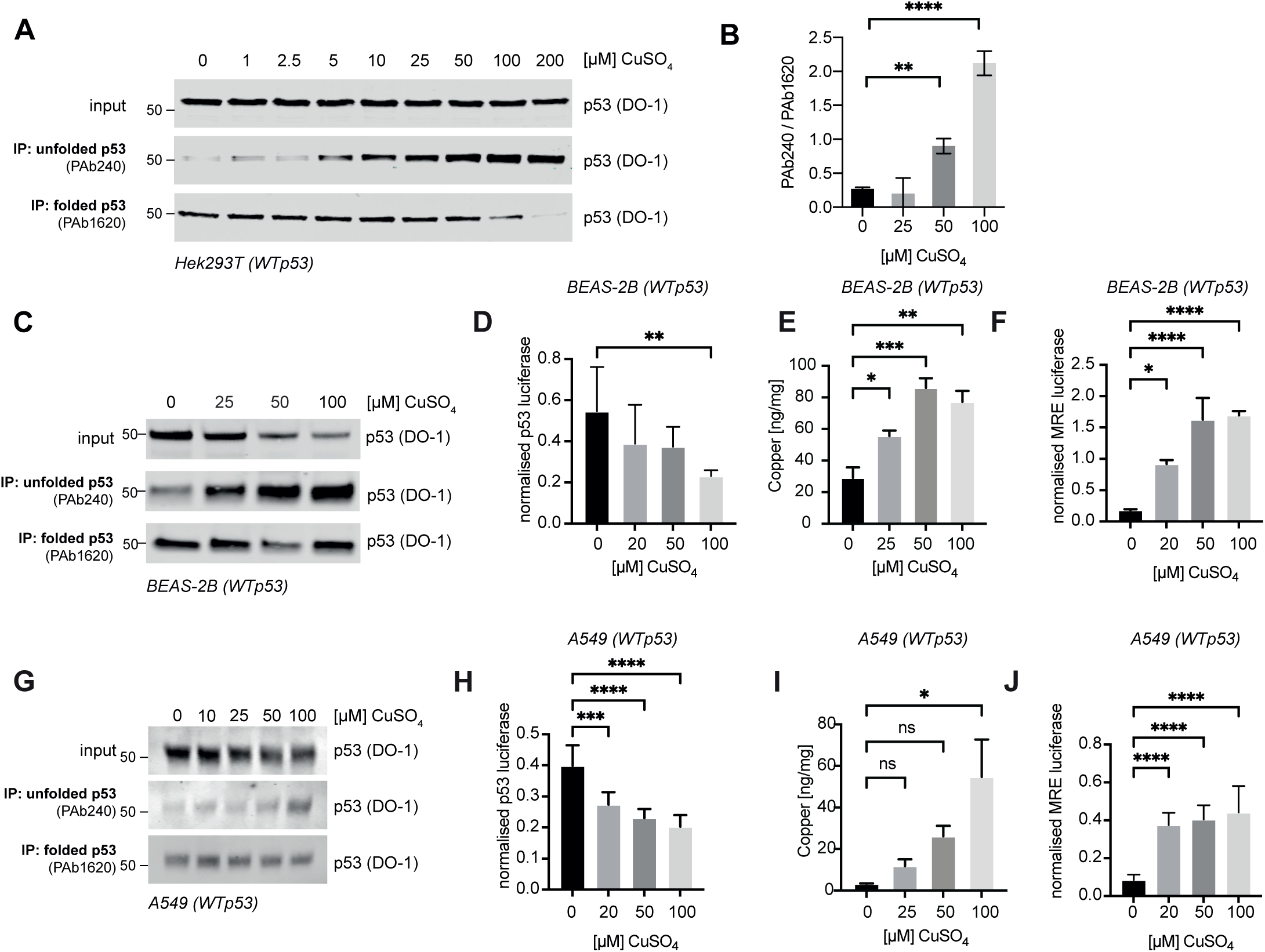
Copper promotes unfolding and inactivation of WTp53. A) Immunoprecipitation of p53 in HEK293T cells (WT p53) using PAb1620 (folded) and PAb240 (unfolded) antibodies, detected by western blot using DO-1. Cells were treated with indicated concentrations of CuSO_4_ for 24h. B) In-Cell western signal intensities of p53 conformational antibodies PAb1620 and PAb240. n=4. See Figure S1A) C) Immunoprecipitation of p53 in Beas-2B cells (WT p53) using the conformation-specific antibodies PAb1620 (folded) and PAb240 (unfolded) and western blot to detect p53 with DO-1 antibody. Cells were treated with indicated concentrations of CuSO_4_ for 24h. D) Beas-2B cells transfected with a p53 luciferase or TK renilla construct and incubated in copper for 24h. Luciferase values were normalised for renilla expression with error bars showing SEM of n=4. E) ICP-MS to determine copper levels in Beas-2B cells exposed to CuSO_4_ for 24h. Metal levels are corrected for protein levels. Error bars showing SEM of n=3. F) Beas-2B cells were transfected with a MRE luciferase or TK renilla construct and incubated in copper for 24h. Luciferase values were normalised for renilla expression with error bars showing SEM of n=4. G) Immunoprecipitation of p53 in A549 cells (WT p53) using the conformation-specific antibodies PAb1620 and PAb240, detected by western blot using the DO-1 antibody. Cells were treated with indicated concentrations of CuSO_4_ for 24h. H) A549 cells were transfected with a p53 BE luciferase or TK renilla construct and incubated in copper for 24h. Luciferase values were normalised for renilla expression with error bars showing SEM of n=3. I) ICP-MS to determine copper levels in A549 cells exposed to CuSO_4_ for 24h. Metal levels are corrected for protein levels. Error bars showing SEM of n=3. J) A549 cells transfected with a MRE luciferase or TK renilla construct and incubated in copper for 24h. Luciferase values were normalised for renilla expression with error bars showing SEM of n=3.

### Copper is accumulated in cancer cells of lung cancer patients, but not in adjacent tissue or serum

As relatively low amounts of copper affected p53 function, we wondered to what extent copper is accumulated in cancers and if such levels would be sufficient to impact p53 function. Possibly the highest risk for copper exposure is through respiratory exposure (e.g. cigarette smoking, vehicle and industrial exhausts), while increased exposure through diet is less of a risk as dietary uptake and release of copper are well regulated through dedicated copper transporters (Focaccio et al., 2024). We therefore speculated lung cancers might be most affected by copper exposure.

Many studies have looked at biomarkers for copper overload and amongst these is mRNA expression of the copper importer *SLC31A1,* which is overexpressed in cancers, and found to be both predictive to disease progression and negatively correlated with patient survival(Kong et al., 2023; Qi et al., 2023; Yun et al., 2022). Using the Xena Functional Genomics Explorer (Xena Browser)(Goldman et al., 2020), we looked at the expression of *SLC31A1* in various cancers and normal tissue using the TCGA GTEx dataset, revealing higher expression in cancers and specifically in lung cancers (figure 2A and 2B, figure S2A). Looking at cancer patient survival, high *SLC31A1* expression correlated to a marginally worse survival in all cancers (TCGA Pan Cancer dataset). When selecting all lung cancers or the more aggressive Squamous Cell Carcinoma in this dataset, higher expression levels of *SLC31A1* also correlated to worse survival, but this was only significant in lung Squamous Cell Carcinoma patients (Figure 2C and D and Figure S2B). Although *SLC31A1* might be correlated to copper levels, it’s expression is considered not directly regulated by copper levels in human cells (Muller et al., 2007). There, the most copper-responsive genes are metallothioneins (Muller et al., 2007), of which metallothionein 2A (*MT2A)* is often highest expressed in cancers and was recognised as a biomarker for non-small cell lung cancer survival(Werynska et al., 2013). Remarkably, in the same datasets as used for *SLC31A1*, *MT2A* expression was lower expressed in all cancer or lung cancers compared to normal tissue (Figure 2E and F) and the differential expression between tumour tissue and controls wasn’t as high in lung cancers as *SLC31A1* (Figure S2B). Despite not finding higher expression in tumour tissue compared to normal tissues, in all cancers, in all lung cancers and specifically in Lung Squamous Cell Cancer, higher expression correlated to a significantly reduced survival (Figure 2G and 2H and Figure S2D). These data suggest copper accumulation in lung cancers and a correlation with reduced survival.

**Figure 2.**
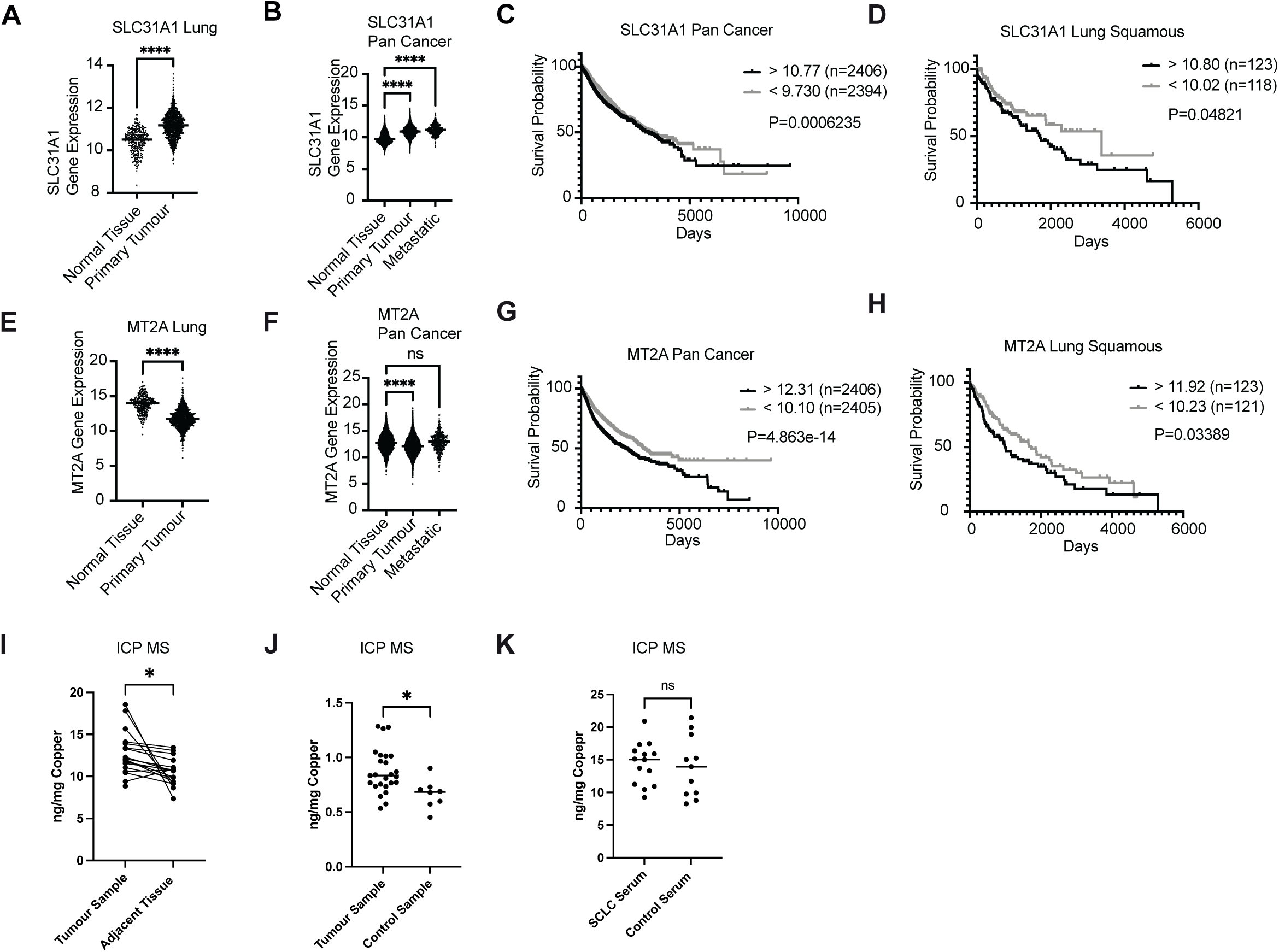
Copper accumulates in cancer cells. A) Median *SLC31A1* expression in normal tissue or primary tumours of the TCGA GTEX database (selected for lung). B) Median *SLC31A1* expression in normal tissue, primary tumours or metastases of all samples in the TCGA GTEX database. C) Overall survival of all cancer patients with low or high *SLC31A1* expression levels using the TCGA Pan Cancer database. D) Overall survival of all cancer patients with low or high *SLC31A1* in the TCGA Squamous lung cancer patient group. E) Median *MT2A* expression in normal tissue or primary tumour of the TCGA GTEX database selected for lung. F) Median *MT2A* expression in normal tissue, primary tumours or metastases of all samples in the TCGA GTEX group. G) Overall survival of all cancer patients with low or high *MT2A* expression levels using the TCGA Pan Cancer database. H) Overall survival of all cancer patients with low or high *MT2A* in the TCGA Squamous lung cancer patient group. I) ICP-MS measurement of copper per tissue content in tumour samples of adjacent tissue of SCLC patients. A pairwise analysis was used to determine statistical significance using a T-test J) ICP-MS measurement of copper per amount of tissue in tumour samples compared to controls. K) ICP-MS measurement of copper per amount of serum in SCLC patients compared to controls.

Notably, *SLC31A1* and *MT2A* are not exclusively regulated by copper. *SLC31A1* is also regulated by other oncogenes, while *MT2A* can be regulated by other metals as well as stress, such as oxidative stress (Cheng et al., 2020; Huo et al., 2023; Ling et al., 2016). To prove that cancer cells indeed have higher levels of copper, we used ICP-MS to measure copper levels in serum, in tumour tissue and tumour adjacent tissue of SCLC patients. As a control, we used patients with suspected lung cancer that had a biopsy taken, but did not have signs of malignancy (Table 1A and B). SCLC status was validated by a histopathologist. In total, we obtained 26 fresh frozen tumour tissues from SCLC patients. Of these, 19 patients had matching adjacent tissue and 14 matching serum (Table 1A and B). As expected, all SCLC patients were smokers or ex-smokers (2 unknowns), which we could match with ex-smokers in the control group (Figure S2E). The average age in the groups (63 versus 68) not significantly different (Figure S2F). Slightly more females were present in the SCLC group than in the control group (Figure S2G). One person in the control group with a hamartoma (a benign tumour that has been linked to Welder’s lung and therefore, perhaps not surprisingly, had the highest levels of copper of all patients in both serum and lung tissue) was excluded (Figure S2H and S2I).

We noted a significant increase in copper levels in tumour tissue compared to adjacent tissue using pairwise analysis in the same patients (Figure 2I). Copper levels were also higher in tumour tissues compared to control samples (Figure 2J). Interestingly, despite previous reports that showed that serum copper levels were predictive for lung cancer mortality, we did not observe differences in the serum of patients compared to controls(Zabłocka-Słowińska et al., 2020) (Figure 2K), suggesting that serum copper levels do not always reflect copper overload in the tumour. Copper levels in tumours or adjacent tissue from smokers or ex-smokers did not differ significantly, suggesting that smoking status did not contribute to differences observed in Figure 2I and J (Figure S2J and S2K). This was further validated by directly comparing tumour SCLC tissue from ex-smokers to normal tissue from ex-smokers, which showed that copper levels were still higher in tumour samples than in adjacent tissue (Figure S2L).

In conclusion, our data demonstrates copper accumulation in cancer cells of lung cancer patients.

### Copper promotes reversible, ROS-independent unfolding of WTp53

We next sought to demonstrate copper-mediated unfolding of p53 in the patient-derived SCLC cell line, CDX10, which is maintained as xenograft in immune compromised mice(Simpson et al., 2020). WT p53 status was confirmed by Sanger Sequencing and p53 could be detected at low expression levels (Figure 3A). Small pieces of xenografted tumours were grown for 48 h ex vivo and exposed to 10 or 100 µM of copper, followed by p53 immunoprecipitation (Figure 3A). These experiments show that p53 in this SCLC cell line exists in a predominantly folded state and that p53 in *ex vivo* SCLC cultures is sensitive to copper-induced unfolding, similar to our findings in other cell lines in 2D plastics (Figure 1A, C and G). Unfolding in response to copper does not seem confined to lung cells as many other cancer cells, including A2780, p53-transfected Hep3B and MCF-7 are sensitive to copper-mediated unfolding of p53 (Figure 3B, Figure S3A and S4A). However, the variation in concentrations needed to unfold p53 with HEK293T cells unfolding at 2.5 µM (Figure 1A) and MCF7 at 200 µM CuSO_4_ is remarkable (Figure S3A). These differences might reflect variations in copper homeostasis between cells, likely due to differences in buffering capacity, copper importer levels or copper exporter levels. To obtain further insights into the dynamics of the copper-induced unfolding of WTp53, we exposed HEK293T cells to increasing CuSO_4_ concentrations for increasing amounts of time. Unfolded WTp53 was already observed after 2 h or 4 h of exposure with 50 µM CuSO_4_, but a decrease in WTp53 conformation was only observed after 24 h of CuSO_4_ treatment (Figure S3B). This 4 h copper-dependent unfolding was readily reversible, as recovery of folding was seen within 4h of copper withdrawal (see immunoprecipitation with PAb1620, 100 or 200 µM CuSO_4_) (Figure S3C). Co-incubation of HEK293T cells with CuSO_4_ and 100 µM ZnSO_4_ prevented some of the unfolding of WTp53 upon CuSO_4_ exposure (Figure S3D).

**Figure 3.**
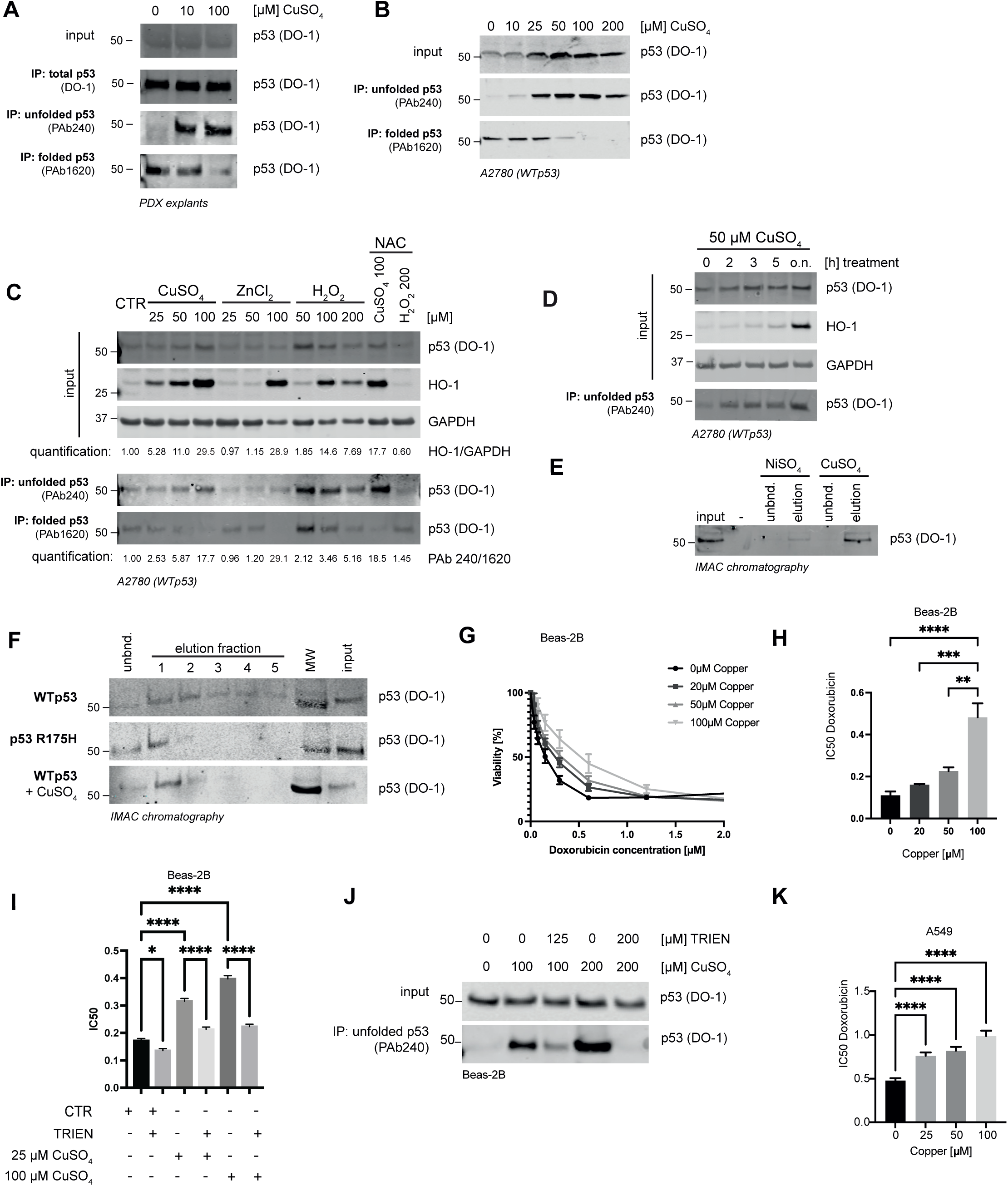
Copper-induced p53 unfolding is not dependent on ROS and leads to doxorubicin resistance. A) Immunoprecipitation of p53 using the conformation-specific antibodies PAb1620 and PAb240 in explants of patient-derived SCLC tumours (WT p53) grown in xenografts. Western blot was used to detect p53 expression using DO-1. Explants were exposed to copper for 24h. B) Immunoprecipitation of p53 using the conformation-specific antibodies PAb1620 and PAb240 in A2780 cells (WT p53) detected by western blot using DO-1. Cells were exposed to copper for 24h. C) Immunoprecipitation of p53 using the conformation-specific antibodies PAb1620 and PAb240 in A2780 cells that were exposed to CuSO_4_, ZnCL_2_, H_2_O_2_, and/ or 100 µM NAC for 24h. p53, GapDH and HO-1 expression levels were determined by western blot. Expression levels were quantified using the Li-Cor. D) Immunoprecipitation of p53 using the conformation-specific antibody PAb240 in A2780 cells that were exposed to CuSO_4_ for indicated times. p53, GapDH and HO-1 expression levels were determined by western blot. E) p53 was in vitro translated and run over a NiSO_4_ or CuSO_4_ IMAC chromatography column. Unbound and elution fractions were run on western blot. F) p53 or mutant p53 R175H was in vitro translated and pre-exposed to 100 µM CuSO_4_ and run over a NiSO_4_ or CuSO_4_ IMAC chromatography column. Input, unbound and elution fractions were run on western blot. MW is molecular weight marker showing the 50kDa band. G) Resazurin survival curves of Beas-2B cells pre-incubated for 24h with CuSO_4_ and then exposed to increasing doses of doxorubicin for 72h. n=4 and error bars indicate SEM H) IC50 of G. n=4. I) IC50 of survival curves of Beas-2B cells (WT p53) pre-exposed to two concentrations of copper and then treated with increasing concentrations of doxorubicin and 125 µM Trien for 72h. J) Beas-2B cells (WT p53) were exposed to copper +/-Trien as indicated and lysates were Immunoprecipitated using the conformation-specific antibody PAb240. P53 was detected using western blot and DO-1 antibody. K) IC50 of resazurin survival curves of A549 cells pre-incubated for 24h with 4 CuSO_4_ concentrations and then exposed to increased doses of doxorubicin for 72h. n=3. Error bars indicate SEM.

Metals including copper are known for their role in the production of reactive oxygen species (ROS) (Cervantes-Cervantes et al., 2005). *In vitro*, p53 is sensitive to unfolding by oxidation, which could be prevented by metal chelators (Hainaut and Milner, 1993a), while ROS were found associated with p53 unfolding in Alzheimer’s disease (Buizza et al., 2012). We therefore wanted to verify if an increase in ROS caused by copper exposure was sufficient to unfold WTp53 in human cells. We verified that ROS induced by H_2_O_2_ (a strong oxidiser) caused unfolding of WTp53 in A2780 cells (Figure 3C). Unfolding was increased by 2-5-fold in the presence of H_2_O_2_ as quantified by dividing the intensity of the bands in the Pab240 by those in the Pab1620 blot (displayed at the bottom of Figure 3C). H_2_O_2_ -mediated unfolding was prevented by co-incubation with the ROS inhibitor N-acetyl cysteine (NAC), reducing unfolding from 5.16-fold to 1.45-fold. (Figure 3C).

Unfolding coincided in all cases with an increase in heme oxygenase (HO-1) expression, which is upregulated by NRF2 upon oxidative stress (Loboda et al., 2016) (Figure 3C). In comparison with copper, zinc seems to only unfold p53 at the high dose of 100 µM. On closer inspection this unfolding seems mostly the result of a marked reduction in the PAb1620 signal (folded) as well as reduced GAPDH levels in the input, rather than an active increase in PAb240 binding. It is therefore unlikely that zinc is causing p53 unfolding directly. When using H_2_O_2_, the PAb240/PAb1620 ratio (unfolded to folded p53) was highest after 100 µM H_2_O_2_ exposure (Figure 3C), but not as high as seen for the highest dose of CuSO_4_. We then used the free radical scavenger NAC at a dose that prevented HO-1 upregulation by H_2_O_2_ and did not cause p53 unfolding (Figure 3C). This dose reduced CuSO_4_-induced HO-1 levels (29.5 to 17.7-fold), but did not prevent copper-mediated p53 unfolding in CuSO_4_-treated cells (17.7 to 18.5-fold). These data suggest that copper-mediated unfolding could be independent of ROS.

If copper had a direct role, p53 folding would be expected to change relatively quickly upon copper exposure. In A2780 cells, p53 unfolding indeed occurred at 2 h after exposure to 50 µM CuSO_4_, at which point HO-1 was not yet upregulated (Figure 3D). These results suggest that changes in copper levels precede transcriptional changes seen upon ROS increase and possibly point to direct effects of copper on p53.

To investigate direct effects of copper on p53 folding we translated p53 in vitro and subjected it to low amounts of copper (Figure S3E). We next used the conformation-specific antibodies for immunoprecipitation and detected a conversion of folded p53 to unfolded p53 upon 5-10 µM of copper exposure (Figure S3E). This prompted us to speculate that copper might bind directly to p53 as has been previously suggested by Hainaut et al, although the copper binding site in p53 is unknown (Hainaut et al., 1995). To determine direct binding, we used Immobilized Metal Affinity Chromatography (IMAC), which has been used by others to detect copper binding to MEK-1 (Turski et al., 2012). We saturated the resin with CuSO_4_ or the nickel salt NiSO_4_. NiSO_4_ was used as a negative control as it is another divalent metal that did not unfold p53 (Figure S3F). WTp53 was successfully enriched in the elution of the CuSO_4_-saturated IMAC resin, while less was detected in the eluates of the NiSO_4_-saturated resin (Figure 3E). To demonstrate specificity, we pre-treated WTp53-containing lysates with CuSO_4_ and used 175H mutant p53 (an unfolded mutant, Figure S3E) which markedly reduced binding of p53 to the resin (Figure 3F). These data could indicate that folded p53, but not an unfolded p53 protein can directly bind copper.

### Copper-mediated p53 unfolding causes chemoresistance

As copper caused p53 unfolding and inhibited p53 transcriptional function, we speculated that it might also cause chemoresistance. We exposed Beas-2B cells to increasing copper concentrations and determined cell viability upon doxorubicin exposure. These studies clearly show a dose-dependent increase in doxorubicin resistance upon copper exposure without affecting cell viability (Figure 3G and H and Figure S3G). Copper-mediated doxorubicin resistance could be reversed by treating the cells with the copper chelator Triethylenetetramine (Trien) (Figure 3I), which also prevented p53 unfolding upon copper exposure (Figure 3J). Copper-mediated chemoresistance could also be observed in A549 cells without affecting cell viability (Figure 3K, Figure S3H).

### Copper causes WTp53 to bind to mutant p53 interactors

Previously, we discovered that WTp53 unfolded by loss of the chaperone CCT, can behave like mutant p53(Trinidad et al., 2013). Many mutant p53 proteins are unfolded and impaired in p53 function(Muller and Vousden, 2013). We therefore wondered whether copper-induced unfolded p53 could also act like a mutant p53 protein. Conformational mutant p53 proteins (for example p53-R175H) can bind p63, or p73 whose functions are suppressed upon this interaction. Notably, WTp53 does normally not bind p63 or p73 (Di Como et al., 1999a; Gaiddon et al., 2001). Binding of mostly folded DNA-contact mutants (for example p53-R273H) to p63 or p73 is possible, but less pronounced compared to conformational mutants(Muller et al., 2009). To elucidate if copper-unfolded WTp53 acquires binding propensity to p53 family members, we used the p53-null line Hep3B to express WTp53 and mutant p53 (R175H and R273H) in combination with TAp63 or TAp73. When transfected with WTp53, p53 unfolds at levels of about 10 µM in Hep3B cells (Figure S4A) and is impaired in transcriptional function in luciferase reporter assays (Figure S4B) upon copper accumulation (Figure S4C). Using Co-immunoprecipitations of transfected HA-tagged TAp63α with p53, we noted that p63 only bound to WTp53 in response to CuSO_4_ (Figure 4A). R175H and R273H mutant p53 bound to p63 in the absence of copper, although this interaction was potentiated in the presence of copper. Similar results were found using TAp73α instead of TAp63α (Figure S4D). The interaction between TAp63α and p53 was verified intracellularly, using PLA, (Figure 4B, Figure S5A). Low levels of copper up to 25 µM increased the PLA signal in the nucleus. Higher concentrations resulted in a signal in the cytoplasm, which suggests copper could promote re-localisation of p63 and mutant p53 to the cytoplasm, which could be the result of apoptotic stress. The increased interaction between TAp63α and p53 coincided with a decreased function of TAp63 as examined using the promoter of the p63 target gene K14 for luciferase reporter assays (Figure 4C), which also showed reduced p63 activity in the presence of 175H mutant p53.

**Figure 4.**
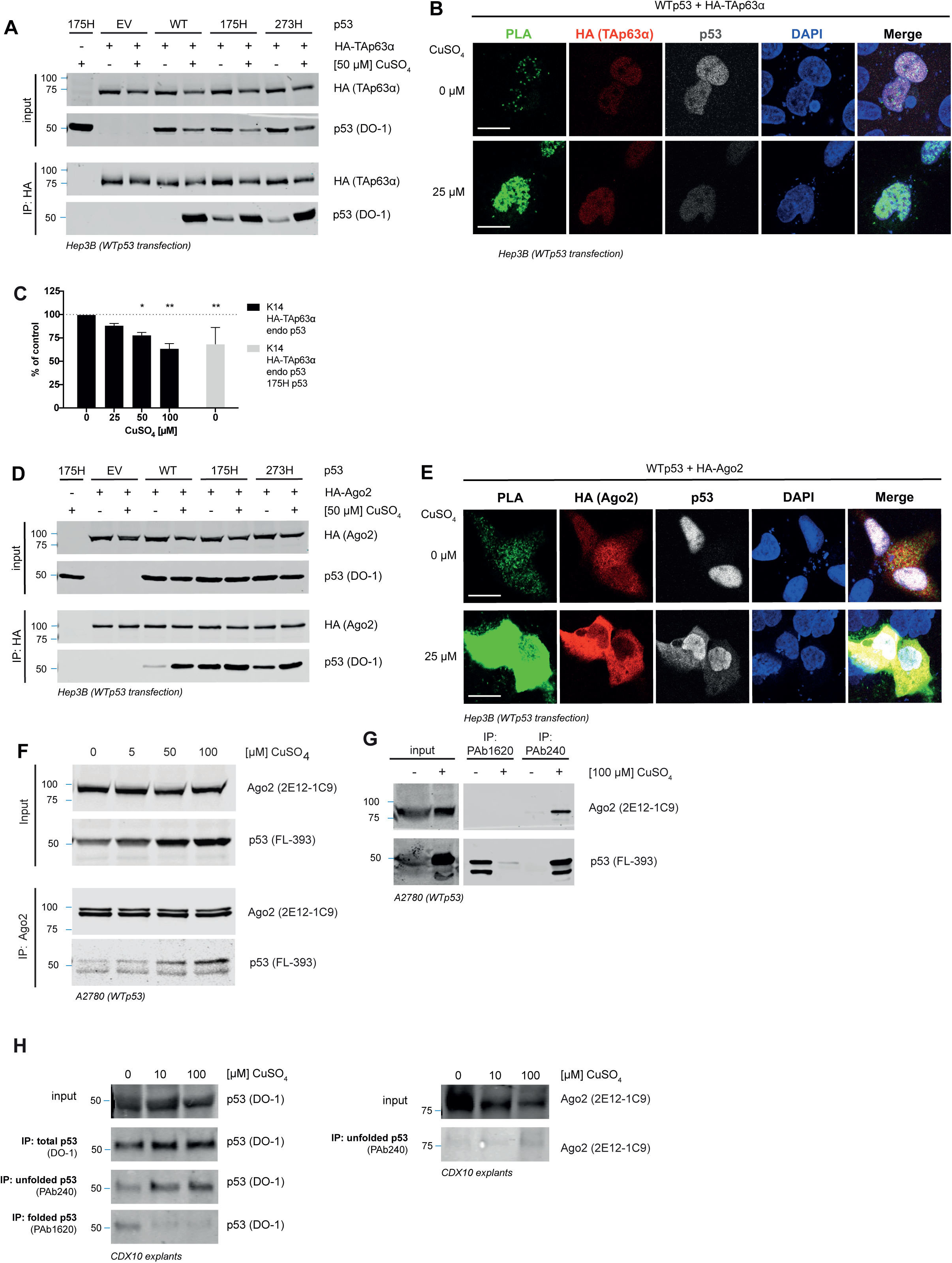
Copper increases the interaction of wt and 273H p53 with TAp63α and Ago2. A) Co-immunoprecipitation of p53 with HA-TAp63α. Hep3B cells were transiently transfected with HA-TAp63α in combination with either empty vector (EV), wt, 175H or 273H p53. Hep3B cells (p53 null) were treated with indicated concentrations of CuSO_4_ for 8h and p53 and HA expression examined using western blot. B) Hep3B cells (p53 null) were transfected with HA-TAp63α in combination with empty vector (EV), wt, 175H or 273H p53. Hep3B cells were treated with 25 µM CuSO_4_ for 8h and fixed for proximity ligation assays. The signal of each individual antibody was assessed after PLA using immune fluorescence. DAPI is shown as nuclear stain (please see Figure S5 for additional conditions and control). Scale bars indicate 20µm C) Luciferase reporter assay of A2780 cells (WT p53) transfected with HA-TAp63α and p63 target gene Keratin 14 promoter-luciferase construct. Cells were treated with CuSO_4_ for 24h. Data are presented as mean ± SEM of three independent experiments. D) Co-immunoprecipitation of p53 with HA-Ago2. Hep3B cells were transiently transfected with HA-Ago2 in combination with either empty vector (EV), wt, 175H or 273H p53. Hep3B cells were treated with indicated concentrations of CuSO_4_ for 8h and p53 and HA expression examined using western blot. E) Hep3B cells were transfected with HA-Ago2 in combination empty vector (EV), wt, 175H or 273H p53. Hep3B cells were treated with 25 µM CuSO_4_ for 8h and fixed for proximity ligation assays. The signal of each individual antibody was assessed after PLA using immune fluorescence. DAPI is shown as nuclear stain (please see Figure S5 for additional conditions and control). Scale bars indicate 20µm F-G) A2780 cells (WT p53) were treated with CuSO_4_ for 24h. Cells were lysed and Ago2 or p53 conformation specific antibodies were used for immunoprecipitation. Endogenous Ago2 or p53 FL353 antibodies were used for detection on interaction partners using western blot. H) SCLC CDX10 ex vivo cultures were treated with CuSO_4_ for 24h and p53 conformation and Ago2 binding determined using immunoprecipitation with conformation antibodies. Ago2 binding or p53 folding were detected using western blot.

Mutant p53 is also known to bind to other proteins and affect their function, such as Argonaute-2 (Ago2) and impact on microRNA biogenesis (Krell et al., 2016). Similar to p63, in the presence of CuSO_4_, Ago2 was able to better bind WTp53 than under normal conditions (Figure 4D). Both R175H and R273H mutant p53 could interact with Ago2 without metal exposure, although copper did slightly increase the interaction. With PLA, we validated this interaction occurred throughout the cell (Figure 4E, Figure S5B). We also verified binding endogenously in A2780 cells and observed an enhanced pulldown of WTp53 by Ago2 at increasing copper concentrations (Figure 4F). This increase could be due to elevated amounts of p53 in the input of this experiment. We therefore used the conformation-specific antibodies to Co-IP Ago2. In the PAb1620 IP (folded p53) in the absence of copper, no Ago2 is detected in the pulldown, indicating that folded p53 did not bind to Ago2. In contrast, in the PAb240 IP (unfolded p53) that pulled down similar levels of p53 as the PAb1620, in the presence of copper a specific pulldown of Ago2 was detected (Figure 4G), showing that unfolded p53 is binding to Ago2 when cells are exposed to copper. An increased amount of Ago2 binding to p53 could also be detected in the SCLC patient CDX ex-vivo cultures that were exposed to copper (Figure 4H).

Together, these data suggest that copper-unfolded WTp53 behaves similarly to mutant p53 by binding to mutant p53 exclusive binding partners.

### Copper causes invasion and tumour growth

To determine if unfolded p53 could promote a ‘mutant p53’ type of GOF in invasion, HEK293T cells were grown into organotypic matrices. We used HEK293T empty vector (EV-Control) and HEK293T cells with a stable p53 knockdown (shRNA) cells or EV-Control cells that express WTp53. These data showed increased invasion into the collagen matrix upon exposure to 5 µM CuSO_4_, whereas HEK293T p53-Knockdown cells did not exhibit an increased invasion (Figure 5A). Knockdown was confirmed by western blot (Figure 5B). To prove that p53 was unfolded in the EV-control cells, HEK293T cells from a parallel matrix exposed to 5 or 10 µM CuSO_4_ were mechanically detached and used for conformational IPs. Unfolding was readily detected at 5 µM (Figure 5C and D), suggesting that unfolded p53 was promoting invasion. To validate the functional consequence of p53 unfolding, we also used A549 cells and examined invasion in organotypic invasion assays into collagen using an air-liquid interface (Figure 5E), and inverted invasion assays where cells invade for 5 days through pores into a matrix containing growth factors (Figure 5F). In both cases, invasion was increased in the presence of copper, at concentrations of around 0.5-1 µM at which p53 did not unfold in 2D after 24 h (Figure 1G). We therefore cultured A549 cells in the presence of CuSO_4_ for 72 h in low serum and could already detect WTp53 unfolding at concentrations as low as 1 µM, suggesting p53 is even more sensitive to unfolding in low serum when subjected to copper (Figure 5H).

**Figure 5.**
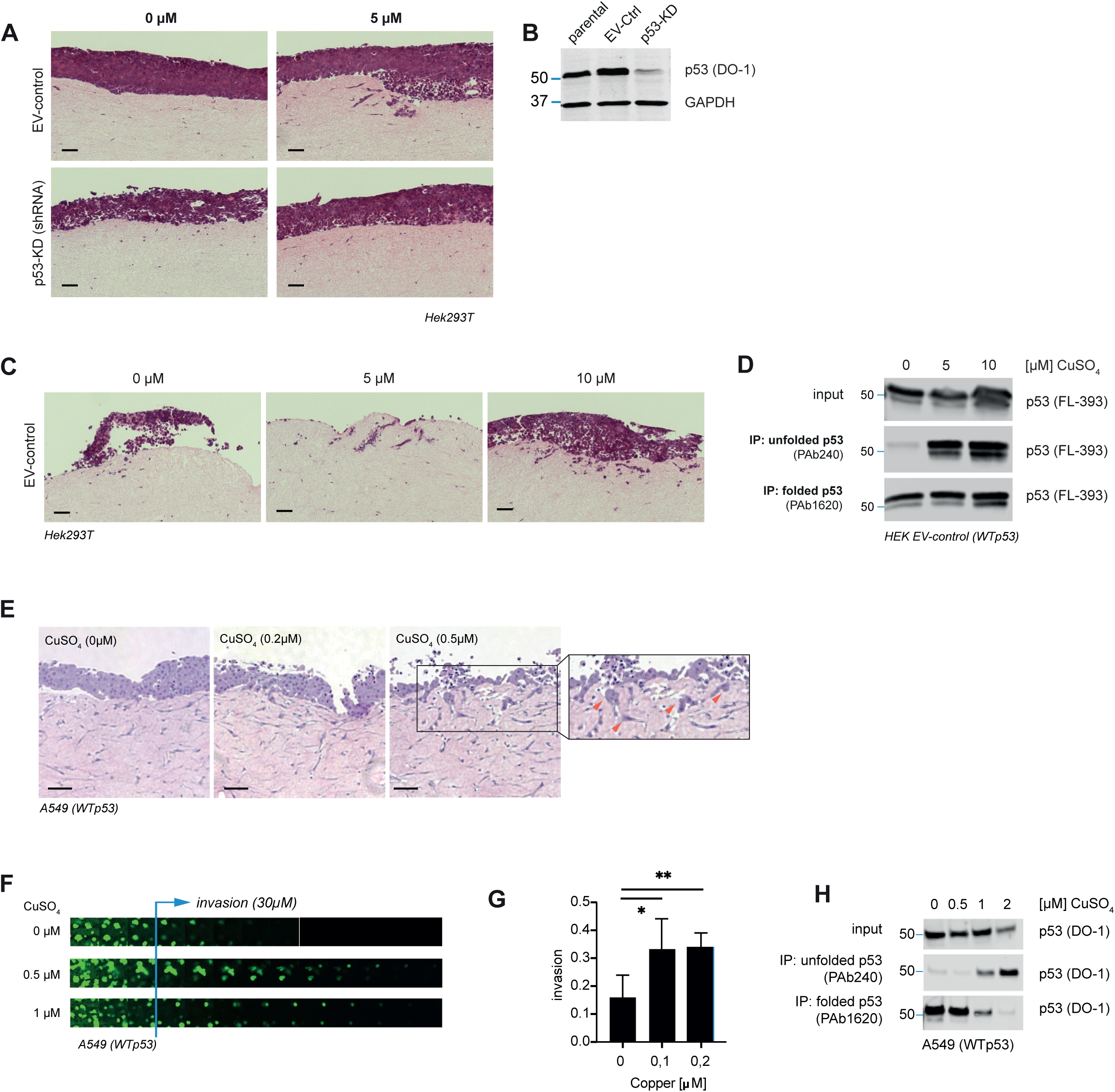
Copper induces invasion. A) HEK293T cells (WT p53) stably transfected with a p53 knockdown construct (p53-KD) or control (EV-ctr) were cultured on fibroblast-contracted collagen matrices, using an air-liquid interface and treated for 14 days with 5 µM CuSO_4_. Invasion was measured in histological sections using H&E staining. Scale bars indicate 50µm B) Western blot validating p53 expression levels of the HEK293T cell lines. GapDH is loading control. C) HEK293T cells were cultured on fibroblast-contracted collagen matrices, using an air-liquid interface and treated for 14 days with CuSO_4_. Invasion was measured in histological sections. Scale bars indicate 50µm D) p53 folding status of cells on the surface of the organotypic assays in C was determined using immunoprecipitation with the conformation antibodies and p53 detection using western blot. E) A549 cells (WT p53) were cultured on fibroblast-contracted collagen matrices, using an air-liquid interface and treated for 14 days with CuSO_4_. Invasion was measured in histological sections. Scale bars indicate 50µm F) A549 cells were placed in inverted invasion assays and invasion beyond 30 µm imaged using confocal microscopy and Z-stacks. G) quantification of invasion beyond 30 µm of F. H). A549 cells were cultured in the absence of medium 2 days prior to copper incubation (in serum free medium). p53 folding was examined using immunoprecipitation with folding antibodies and p53 expression measured using western blot.

### Certain mutants are more vulnerable to copper accumulation and such mutations are found frequently in lung cancers

In more than 50% of cancers, p53 is mutated resulting in the expression of a mutant protein. In many cases this mutant p53 is already unfolded, or non-functional, but partially-functional mutants exist. Such partially-functional mutants can still exert WT activity but might be more prone to unfolding in response to copper than WTp53. According to The TP53 database (de Andrade et al., 2022), 313 missense mutations that occur within p53 render the molecule partially functional. These occur in 2.43% of all primary tumours (221/9104, TCGA pan-cancer dataset) and, interestingly, are doubly frequent in lung cancers, occurring in 5.24% of all primary samples (52/993, TCGA lung cancer) (Figure S6). Partially functional and functional missense mutations are mostly found in exon 5 of p53 (Figure S6). We therefore speculated that in lung cancers with partially functional mutations, the increased copper levels could be inactivating these mutants fully, causing more chemoresistance and more invasion or metastasis.

We selected a partially-functional mutant A159S, that occurs in lung cancer and examined its p53 function. The p53 target genes *BBC3* (Puma), *C12ORF15* (TIGAR) and *CDKN1A* (p21) were similarly expressed upon overexpression of WTp53 or mutant p53 A159S (Figure 6A-D). Importantly, mutant p53 A159S and WTp53 expressed at similar protein levels (Figure 6E). The p53 target gene *BAX* was slightly higher expressed upon WTp53 expression than upon mutant A159S expression. As a negative control, the non-functional and conformation mutant R175H was unable to induce any of the genes. Upon a short period of 6 hr of copper stimulation, WTp53 mediated expression of *BBC3* and *CDKN1A* was slightly decreased, but expression of *BBC3*, *CDKN1A* and *C12ORF15* was much lower in cells expressing the A159S mutant under these conditions. This is despite the fact that copper marginally increased the expression of *BBC3* and *CDKN1A* in R175H and control transfected cells (Figure 6A and C). Interestingly, copper did not seem to affect *BAX* or *C12Orf5* when p53 wasn’t expressed or R175H was transfected. Protein expression of *CDKN1A* (p21) was also examined by western blot and showed a more profound reduction in expression upon copper exposure in A159S transfected cells than in WTp53 transfected cells (Figure 6E). These data suggest copper profoundly prevents mutant p53 A159S from acting as a p53 transcription factor and that there might be specificity in which p53 target genes are inhibited by copper.

**Figure 6.**
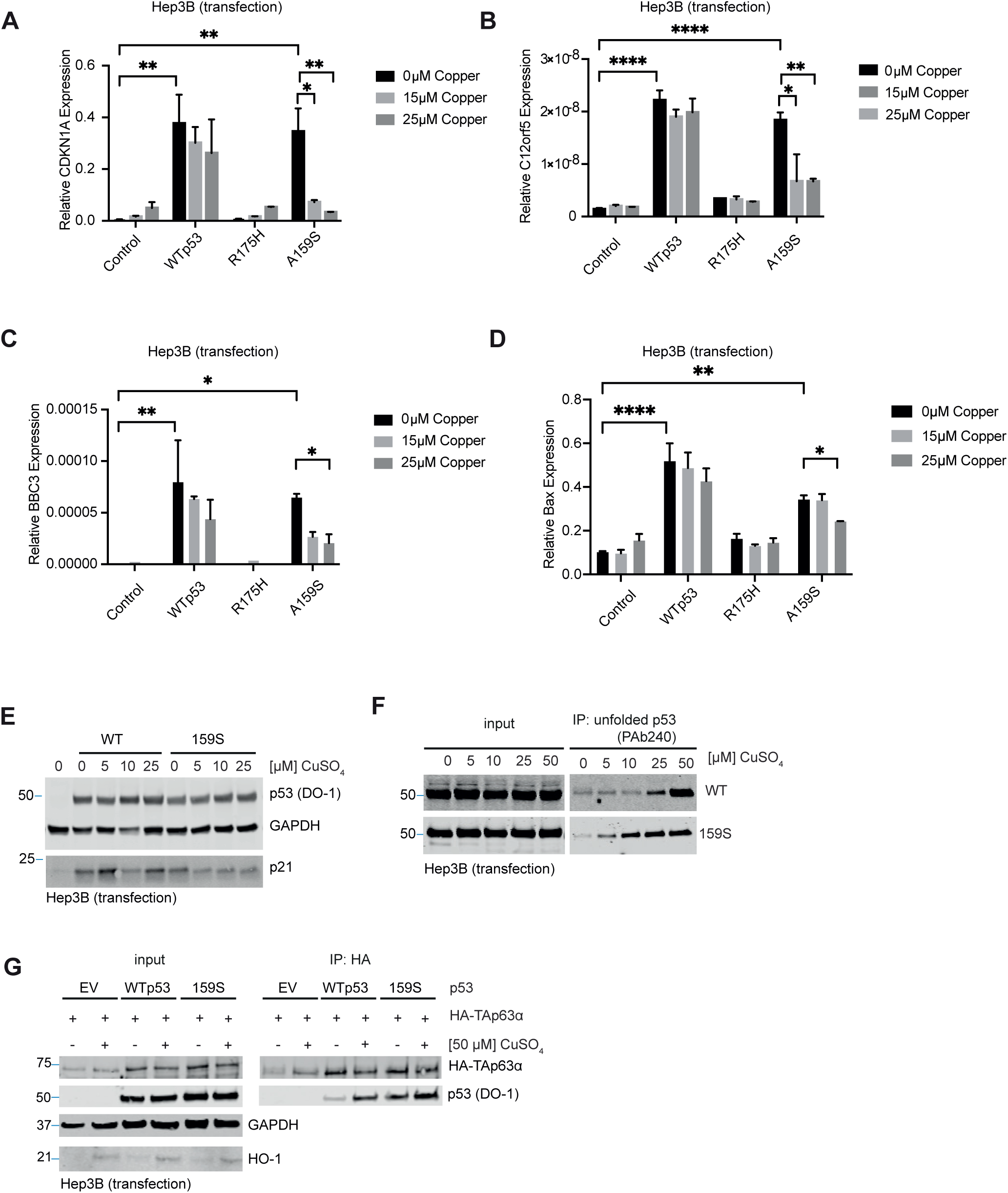
The partially-functional p53 mutant A159S is more affected by copper than WTp53. A-D) Hep3B cells (p53 null) were transfected with EV, WT, 175H or 159S mutant p53 and subjected to 0,15 or 50 µm CuSO_4_. Expression of *CDKN1A, C12ORF5, BBC3* or *Bax* were measured using qRT PCR with GapDH as reference gene. N=3. E) Hep3B cells were transfected with EV, WT or A159S mutant p53 and subjected to 0, 5, 10 or 25 µm CuSO_4_ after which p53, p21 or GAPDH levels were examined on western blot. F) Hep3B cells were transfected with WT or A159S mutant p53 and subjected to 0, 5, 10, 25 or 50 µm CuSO_4_, after which p53 folding was analysed using co-immunoprecipitation with the 240 antibody. G) Hep3B cells were transfected with EV, WT or 159S mutant p53 in conjunction with HA-TAp63 and subjected to 50 µm CuSO_4_, after which HA was immunoprecipitated and p53 binding determined on western blot. GapDH was used as loading control and HO-1 is shown in the input as a measure of low copper load.

Importantly, 5 µM of copper already resulted in unfolding of A159S mutant p53, whereas unfolding in WTp53 expressing cells could only be detected at concentrations of 25 µM after 6 h of copper incubation (Figure 6F). Importantly, the A159S mutant has a greater ability to bind to HA-p63 under normal concentrations, which is further enhanced in the presence of 50 µM of CuSO_4_ (Figure 6G). These data suggest that A159S mutant p53 is more impaired in WT function by copper than WTp53, despite being able to induce certain p53 target genes to similar extents under normal conditions. These data also show that 159S’s GOF is exacerbated. Together these data suggest that partially functional mutants such as 159S are more vulnerable to copper increases than WTp53.

## Discussion

Many studies have focussed on the role of copper on cancer cell metabolism (Ge et al., 2022). Copper is used by cancer cells to proliferate and to adjust extracellular matrix to aid invasion into other tissues. For this reason, chelation strategies have been considered as a treatment regime in cancers. These include, but are not limited to penicillamine, Trien, TTM and disulfiram (Bian et al., 2023). Until now results of such trials have been variable (Tang et al., 2023), suggesting that we do not fully understand the role of copper in cancers.

In our manuscript, we treated Beas-2B cells with the copper chelator Trien and noticed that chemoresistance and p53 unfolding could be avoided or reversed using this chelator alongside copper exposure. Trien has been explored as a copper-lowering drug in clinical trials, given alongside platinum drugs (Fu et al., 2014; Fu et al., 2012; Kuo et al., 2012), but is not yet routinely used in the clinic. Based on our study, perhaps p53 status needs to be considered when selecting whom to treat with chelators. From our results, only patients with WTp53 or a mutant p53 that has some residual p53 activity and likely capable of refolding (listed as partially functional mutants in the p53 database) could benefit from these treatments.

Many studies have focused on refolding mutant p53 using small molecules that can act by facilitating zinc donation to mutant p53, in order to restore at least some original/physiological/WT folding and concomitant WT function (Blanden et al., 2015). Despite this focus on mutant p53, WTp53 itself is already unstable and can exist in a folded or unfolded state under normal conditions (Bell et al., 2002; Cañadillas et al., 2006).

Copper was found to directly affect WTp53 folding at concentrations that were non-toxic. In this manuscript, we demonstrate that small increases in copper as seen in small cell lung cancer patient cells can impact p53 by unfolding the protein, likely directly by binding to p53. Unfolding of p53 impairs its normal WT function, but can also promote a GOF that manifests as increased invasion and chemoresistance. This means that small increases in copper not only cause loss of tumour suppression, but may also potentiate aggressiveness of the tumour.

Remarkably little is known about the actual copper levels within tumour cells. Actual copper levels are frequently not measured and instead association studies with copper homeostasis proteins or gene expression are used. Using such biomarkers from publicly available databases, we saw that *SLC31A1* expression was raised in cancer tissue and higher expression levels correlated with a worse survival. Another biomarker is metallothionein, which is upregulated in response to copper. Although increased expression correlated to survival, expression levels of *MT2A* were actually lower in tumour tissue than in control tissue. Other biomarkers that are often used in literature are copper levels in blood, serum or urine. Few studies have shown copper levels in cancers and to our knowledge only one study reported lung cancers to have higher copper levels than control samples, but this study predominantly consisted of lung adenocarcinoma patients (Díez et al., 1989). In our study, we can see an increased level of copper within the cancer cells of small cell lung cancer patients. Copper levels were increased compared to lung tissue in controls, but also compared to adjacent tissue of the same patient. Remarkably, serum levels did not reflect the increased copper levels in the lung cells and suggest that serum copper levels might not be a good biomarker for copper accumulation in the lung. Ideally, we identify a biomarker that not only shows copper accumulation, but also p53 function and folding status. Folding status of p53 is difficult to see in patient samples as freezing or fixation techniques destroy the native folding state of the protein and do not allow for detection using the antibodies used in this study. We are currently using mass spectrometry techniques such as LIP-MS (limited proteolysis Mass spectrometry) to identify easily accessible peptides that can be detected in folded or unfolded p53. Alternatively, perhaps we can identify conformation-specific p53 target genes. Such biomarkers would help us select those patients that could be treated with a copper chelator.

More recently, the focus of cancer researchers has shifted to another intriguing aspect of copper homeostasis, cuproptosis; a type of cell death that results from excessive amounts of copper affecting metabolism genes(Tang et al., 2024). This copper binds to lipoylated proteins involved in mitochondrial energy processes that then aggregate and cause proteotoxic stress. Although a direct role of p53 in cuproptosis remains elusive, the fact that p53 inhibits glycolysis, enhances mitochondrial respiration and regulates glutathione expression suggest it at least indirectly primes cells for cuproptosis (Xiong et al., 2023). As mutant p53 is promoting glycolysis and inhibits mitochondrial respiration, it is likely mutant p53 expression might help cancer cells to resist cuproptosis. Intracellular copper levels can be enhanced by regulating copper homeostasis. An example is ionophores that raise intracellular copper levels (Zhang et al., 2024). Our work suggests that low levels of copper, that would not be sufficient to induce cuproptosis, could inactivate p53 through unfolding. It would be interesting to determine if high levels of copper cause a more acute response and initiate a p53-mediated cuproptosis. This would not be dissimilar to how p53 regulates cell death in response to excessive DNA damage, while triggering cell cycle arrest and repair in response to a low amount of DNA damage.

In terms of exposure to copper and a correlation to cancer formation, copper is currently not considered a carcinogen by the work health organisation (www.apps.who.int/ cas 7440-50-8). As we require copper for many of our essential enzymes it is hard to correlate exposure to a biomarker such as copper deposition in a certain organ, blood, urine etc to cancer risk. This will also depend on the route of exposure, and we don’t know if copper inhalation leads to a similar distribution of copper than dietary copper exposure. Our study shows a discrepancy between copper accumulation in the actual lung tissue and serum levels asnd copper was not seen accumulated in adjacent tissue of lung cancer patients compared to lungs of normal individuals. These data might suggest that copper didn’t initiate the tumour formation, but perhaps helps the tumours propagate. In order to make clear conclusions between exposure and health effects, better biomarkers are needed as discussed in (Bevan and Levy, 2024). In the future, we would like to explore p53 transcriptional function in the presence of copper and potentially use such genes as biomarkers for copper status in cancers. As for exposure, a better animal model that captures inhalation risks associated with copper inhalation might be needed to identify risk of copper exposure on cancer formation, progression and chemoresistance.

## Acknowledgements

We would like to thank Callum Hall for valuable discussions and experimental help, as well as the histology departments of the MRC Toxicology Unit and CRUK Manchester Institute for histological processing of the organotypic matrices. This work was funded by EPSRC EP/S022791/1, Core CRUK Manchester Institute (grant number A27412), Medical Research Council, Toxicology Unit Core funds and by Pleco Therapeutics. CDX work was funded through Core Cancer Research UK (CRUK) funding to the CRUK Manchester Institute (grant number A27412), CRUK National Biomarker Centre (CTRNBC-2022/100001), and the CRUK Lung Cancer Centre of Excellence (grant number A29420). Patient recruitment was supported by the National Institute for Healthcare Research (NIHR) Manchester Biomedical Research Centre, the NIHR Manchester Clinical Research Facility at The Christie Hospital and the CRUK Lung Cancer Centre of Excellence. Pleco Therapeutics funded work regarding the Biobank sample collection.

## Author contributions

YvG and LA designed and executed experiments. HR, SB, ZN and EM executed experiments. MG directed CDX10 expansion and tumour dissection. KS carried out genomic analysis of CDX10.YvG and LA wrote the manuscript, with comments from PAJM. SB and HR corrected the manuscript JLQ classified SCLC patients. PAJM directed the study and acquired funding

All authors approved the manuscript.

## Declaration of AI

No form of AI was used to generate this article

## Material and Methods

### Patients

Frozen SCLC patient and age-matched control lung tissues were obtained from the Manchester Cancer Research Centre (MCRC) Biobank. For some, patients matched serum samples were also available. Details of the patients are listed in table 1A and B. The MCRC Biobank holds a generic ethics approval which can confer this approval to users of banked samples via the MCRC Biobank Access Policy under the Biobank Ethics Code: 18/NW/0092, application code 21/PAMU/1.

### CDX SCLC patient material

The SCLC CDX10 patient cell line was obtained from Prof C.Dive. CDX10 generation was performed as described previously (Hodgkinson et al., 2014; Simpson et al., 2020). CDX models were generated from patients CTCs enriched from blood samples at pre-chemotherapy baseline and/or at post-treatment disease progression time-points (designated P, or PP) (Hodgkinson et al., 2014; Simpson et al., 2020). WTp53 status was confirmed using Sanger sequencing (Hodgkinson et al., 2014).

For ex vivo culture, explants were placed on platinum grids in the presence of DMEM + 10% FBS such as to form an air-liquid interface, and grown at 37°C in 5% CO_2_ atmosphere. After 24h, the explants were diced further using a ceramic blade, and homogenised in NP40 lysis buffer (as described previously) using mechanical shearing with 5 mm stainless steel metal beads (Qiagen). Debris were cleared by centrifugation. Immunoprecipitation was performed as described above.

### Xena Browser and the p53 database analysis

Xenabrowser was used (Goldman et al., 2020) on the TCGA TARGET GTEx dataset to assess the gene expression of *SLC31A1* and *MT2A* in cancer patients compared to controls. Samples without data were excluded. Data from the TCGA Pan-Cancer and the TCGA Lung Squamous Cell Carcinoma dataset was extracted to generate Kaplan Meyer survival curves using Graphpad Prism using only primary tumours. Statistical significance for the violin plots was determined using a one-way ANOVA with multiple-testing correction. Log-rank test was used to determine differences in survival. P53 mutation frequencies were ascertained by extracting a list of all partially inactivating mutations in *TP53*, from The *TP53* Database (R20, July 2019)(de Andrade et al., 2022), and using this as a search prompt in Xena Browser(Goldman et al., 2020) to identify all patients in the TCGA lung cancer and TCGA pan-cancer datasets harbouring such mutations. Frequencies of partially functional mutations, among all primary tumour samples with data on *TP53* mutational status, were then plotted in Microsoft Excel for the respective datasets.

### Cell culture and reagents

HEK293T (origin: kidney, female, ethnicity unknown), MCF-7 (origin: pleural effusion from breast adenocarcinoma, Caucasian female), Hep3B (origin: liver cancer, Black male), Beas2B (origin: lung non-cancerous bronchial epithelium, male, ethnicity unknown) and A549 (lung adenocarcinoma, Caucasian male) cells were obtained from ATCC and cultured in DMEM (Thermo Fisher), supplemented with 10% FBS (Sigma) and 1% Pen/Strep (ThermoFisher) at 37 °Channa. A2780 cells were obtained from Sigma and cultured in RPMI 1640 (Thermo Fisher) supplemented with 10% FBS and 1% Pen Strep. Cells were authenticated using Eurofins and tested for mycoplasm. HEK293T stably knocked down for p53 and corresponding control cells (plasmids pSuper p53 siRNA or pSuper control siRNA) were generated by clonal selection with puromycin (Sigma).

### Plasmids, sequences and transfections

Expression plasmids for WTp53, 175H p53, 273H p53, HA-TAp63α, Keratin-14-luciferase and TK renilla were described before (Muller et al., 2009) . A159S mutant p53 was generated using a KOD hotstart kit (Merck) with the following Eurofins oligos: 159S FW-5’ -CCC CCG CCC GCC ACC CGC GTC CGC TCC ATG GCC ATC TAC AAG CAG -3’ and 159S RV-5’ – CTG CTT GTA GAT GGC CAT GGA GCG GAC GCG GGT GCC GGG CGG GGG -3’. PG13 (WT p53) luciferase was purchased from Addgene and was originally described in (el-Deiry et al., 1993). MRE luciferase was a gift from Dr Klomp (van den Berghe et al., 2007). HA-Ago2 was kindly provided by Dr. Martin Bushel. The pSuper puro vector was purchased from oligoengine and a p53 target sequence (fw: GACUCCAGUGGUAAUCUACUU and rev: UUCUGAGGUCACCAUUAGAUG) was cloned into the vector as per manufacturer’s instructions. For stable knock down of p53, transfection of plasmids was performed using GeneJuice (Merck-Millipore) according to the manufacturer’s protocol. For all other transfections, 1mg/ml Polyethylenimine (PEI 25K, Polysciences) with Opti-MEM (Thermo Fisher) was used in which 1 µg plasmid was combined with 3 µl PEI in 250 µl Opti-MEM, vortexed and inbated for 10 minutes prior to adding dropwise to cells (70% confluent in 6-well).

### Immunoprecipitation and western blot

Cells were lysed in NP-40 lysis buffer [100 mM Tris-HCl pH 8 (Sigma), 100 mM NaCl (Sigma), 1% NP-40 (Sigma)] supplemented with a complete protease inhibitor cocktail (Roche). After 10 min incubation on ice, cell debris was removed by max centrifugation at 4°C for 10 min and input taken. For the IPs, Dynabeads sheep anti-mouse IgG were washed twice with PBS, once with 5% BSA (Sigma), twice with PBS and then pre-coupled tumbling at 4°C for 2 h with antibodies anti-p53 PAb1620 (Merck-Millipore, 0.2 µg/15 µl beads/IP), anti-p53 PAb240 (Merck-Calbiochem, 0.6 µg/15 µl beads per IP), , anti-HA (16B12, Biolegends, 2.25 µg/60 µl beads/IP) or anti-Ago2 (2E12-1C9, Abcam, 5 µg/60 µl beads/IP). Lysates were added to the beads and incubated overnight at 4°C. Beads were washed three times with NP-40 lysis buffer, eluted with 2x sample reducing buffer and boiled for 10 min at 99°C. Input and IP samples were separated in SDS-PAGE and transferred onto nitrocellulose membranes. After blocking with 5% skimmed milk (Marvel) in TBS (Santa Cruz)/ 0.1% Tween20 (Sigma) for 30 min at RT, the membranes were incubated with primary antibodies in 5% skimmed milk/TBS-T. The following primary antibodies were used for western blotting: anti-p53 (FL393 detecting WT p53, Santa Cruz, 0.2 µg/ml or DO-1, Santa Cruz, 0.2 µg/ml), anti-Dicer (H-212, Santa Cruz, 0.8 µg/ml), anti-c-myc (9E10, Santa Cruz, 0.4 µg/ml), anti-HA (16B12, Biolegends, 1 µg/ml) and anti-Ago2 (2E12-1C9, Abcam, 0.83 µg/ml). The blots were washed three times with TBS-T and then incubated with corresponding IRDye secondary antibodies 680RD or 800CW (Li-Cor, 0.05 µg/ml). After three times washing with TBS-T protein signals were visualized using the ODYSSEY Sa infrared system (Li-Cor, Cambridge, UK) and Image Studio software 2.0 (Li-Cor). For the detection of IP Ago2 with truncated mutant p53 constructs, HRP-conjugated light chain antibodies were used with ECL-based detection on film.

### In-Cell Western assay

Cells were seeded in 96 well plates (Greiner, 10.000 per well) and transfected with the different p53 plasmids or incubated in copper as indicated. Plates for HEK293T cells were coated with Poly-L-Lysine (Sigma). Cells were fixed with glutaraldehyde as described by Haim et al. (Haim et al., 2013). The cells were permeabilized and blocked by incubation with 0.2% Triton-X-100 (Sigma) in Odyssey blocking solution (Li-Cor) for 1 h at RT. After three washes with PBS, wells were incubated with antibodies PAb1620 (83 ng/well) or PAb240 (12.5 ng/well) for 2 h at RT. The wells were washed three times with PBS and then IRDye anti-mouse secondary antibody 800CW (Li-Cor, 50 ng/well) was added and incubated for 1 h in the dark. Wells were washed with PBS three times and incubated with anti-p53 FL393 antibody for 2 h at RT. After three washes with PBS, all wells were incubated with IRDye anti-rabbit secondary antibody 680RD (Li-Cor, 50 ng/well) for 1 h and again washed three times with PBS before detecting and quantifying the signals of each well with the ODYSSEY Sa infrared system (Li-Cor) and Image Studio software 2.0 (Li-Cor).

### In vitro translation and copper binding on IMAC columns

*In vitro* translation of p53 was performed in TnT T7 quick-coupled transcription/translation system according to the manufacturer’s protocol (Promega, Southampton, UK). After completed translation, 10 µl of produced proteins were incubated with indicated concentration of CuSO_4_ in a total volume of 50 µl NP-40 for 30 min at RT and then subjected to p53 conformational IPs or bound to IMAC columns.

For metal binding experiments, hiTrap IMAC High performance columns (GE healthcare) were loaded with CuSO_4_, NiSO_4_ or ZnCl_2_. Columns were washed with distilled water (5 column volumes, CV), loaded with 2 ml of 0.1 M CuSO_4_, NiSO_4_ or ZnCl_2_ and then washed with 5 CV of water and 10 CV of binding buffer (20 mM sodium phosphate, 0.5 M NaCl, 0.5 mM imidazole). In vitro translated proteins were diluted to 1 ml in binding buffer and applied to the column. The column was washed with 15 CV binding buffer and eluted with 7 CV of elution buffer (20 mM sodium phosphate, 0.5 M NaCl, 5 mM imidazole). The last 5 CV of the binding buffer and the first CV of the elution were used to constitute the unbound fraction. The last 6 CV of elution were combined as the metal bound fraction of protein. These two 6 ml solutions were concentrated to 100 µl using amicon ultra-4, ultracel-3 (>3 kDa cutoff) concentrators (Millipore). 30 µl of concentrate was loaded for western blot analysis.

### Luciferase assay

A2780 cells were seeded in 24 wells (50.000 per well) and transfected with 300 ng Keratin-14-luciferase construct, 3 ng Renilla luciferase and 200 ng HA-Tap63α in combination with 400 ng EV or 175H p53 constructs, followed by CuSO_4_ incubations. A549 and Beas-2B cells were seeded in 6-well plates and transfected with PG13 or MRE luciferase (600 ng) in combination with 60 ng TK renilla. After 24 h cells were split into 96 well-plates and incubated in increasing CuSO_4_ concentrations. Cells were washed with PBS and then harvested with passive lysis buffer (Promega) and luminescence was measured according to manufacturer’s protocol using the Dual Luciferase Reporter Assay system (Promega) and a GloMax 96 dual injector Microplate Luminometer (Promega, Southampton, UK). Firefly luciferase measurements were corrected for Renilla luciferase values and expressed as percentage of untreated cells.

### Immunofluorescence and Proximity ligation assay (PLA)

Hep3B cells were seeded on glass coverslips and transfected with 0.3 µg HA-TAp63α, HA-Ago2 or EV in combination with 0.3 µg EV, WT or 175H p53 plasmids. After 24 h, cells were treated with CuSO_4_, then washed three times with ice-cold PBS and fixed with 4% PFA for 15 min at 4°C. After fixing, cells were permeabilized with 0.3% Triton-X-100 in PBS for 10 min at RT and then the proximity ligation assay protocol was performed according to the manufacturer’s instructions (Duolink, Sigma). The following primary antibodies were used: anti-HA (1:250, 16B12) and anti-p53 (1:200, FL393). In addition to the PLA signal, the cells were stained for the overexpressed proteins (HA-TAp63α, p53, HA-Ago2) by first washing with buffer A for 2 times 10 min and then incubating with the following secondary antibodies: anti-mouse Alexa Fluor 546 (Life Technologies) and anti-rabbit Alexa Fluor 633 (Life Technologies) for 1 h at 37°C. After washing two times with buffer A, one time with buffer B and one more time with 100 times diluted buffer B coverslips were mounted on glass slides using mounting fluid containing DAPI (Duolink). Representative fluorescent images were obtained using a Zeiss LSM 150 confocal microscope (Zeiss, Cambridge, UK).

### RNA isolation, cDNA synthesis and RT-PCR

RNA was extracted from 10cm^2^ wells using the Total RNA Purification Kit (Norgen Biotek). cDNA was synthesized according to manufacturer’s instructions using the High-Capacity cDNA Reverse Transcription Kit (Applied Biosystems) with 1 µg RNA in a final volume of 20 µl. RT-qPCR was prepared according to manufacturer’s instructions using the PowerUp SYBR Green Master Mix (Applied Biosystems) with 0.5 µM primers and run in the QuantStudio 3 Real-Time PCR System (Thermo Scientific). The used primers are listed in Table S2. Cycling conditions: 40 cycles, Denaturation (95°C, 15s), annealing+extension (60°C, 60s). Melt curve unimodality and negative controls (no RT, no RNA) were confirmed.

### Invasion assays

Invasion in organotypic assays was performed as previously described by Timpson et al. using TIFs to contract a collagen matrix (Timpson et al., 2011). 40,000 cells were seeded on top of the matrix and allowed to become confluent for 4-5 days. Matrices were then transferred on platinum grid scaffolds. The cells were treated for 14 days with CuSO_4_ by replacing the medium every 2-3 days. Matrices were fixed in 4% PFA (Thermo Scientific) for 24h at RT, embedded in paraffin, cut and stained with haematoxylin and eosin. Representative images of invading cells were taken with the Zeiss Axiovert 200M microscope.

The Inverted transwell assay was performed as described previously with the exception of using Geltrex (Muller et al., 2009). Indicated amounts of CuSO_4_ were mixed into a 50/50 solution of PBS and Geltrex (Gibco) and allowed to solidify in transwell chambers (60 µl). A total of 30,000 A549 cells were placed on the bottom of the (inverted) chamber and allowed to settle for 4h. Cells (bottom chamber) were cultured in serum-free DMEM with indicated CuSO_4_ concentrations, while the upper chamber contained DMEM with 10% FCS, 10 ng/ml EGF (Sigma), 10 ng/ml HGF (Sigma), and indicated CuSO_4_ amounts.

Cells were incubated for 5 days, and both chambers were incubated with 4 nM Calcein AM for 1h. Calcein-stained cells in transwells were imaged every 10µm (Z-axis) with an LSM 880 confocal microscope (Zeiss). Occupied cell surface from each layer was quantified using Image J (“Median” filter to smoothen surfaces, “Threshold” to delineate cells from background, “Analyze Particles” to quantify surface). Surfaces starting from the 4^th^ layer (after 30nm) were summed, and divided by the sum of all acquired surfaces to generate an invasion score

### ICP MS

For ICP-MS, cells were grown in 10cm plates and harvested at approximately 80% confluence. Cells were washed 2X with PBS and harvested in 150 µl NP40 lysis buffer and incubated for 15 min RT. 50 µl was used for protein detection using a Pierce BCA assay (Thermo Fisher Scientific) according to the manufacturer’s protocol. The remaining lysate was freeze-dried using a speedvac (Eppendorf 5301) and dissolved in 100 µl HNO_3_ (Emplura 65%), heated to 65 °C for 30 min, followed by boiling at 99 °C for 30 min. For tissues, 50mg of tissue was freeze-dried and dissolved in 100 µl HNO_3_. . Samples are subsequently diluted 50-fold with 2.5% HNO3 (Fisher), containing 10 mg/ ml internal standards (In, Ag, Be), and copper measured by ICP-MS (ThermoFisher iCAP RQ , QCell CRC) in KED mode using pure He gas. Copper standards (0-500 mg/ml matrix matched 2.5% HNO3 + 10 mg/ml) were analysed along with the samples to calculate the metal concentrations using the standard curve.

### Cell viability assay (resazurin reduction assay)

A549 or Beas-2B cells were seeded in 96 well plates (1000 cells/well). 24 h after seeding, cells were incubated with 0, 25, 50, 100 µM CuSO_4_ for another 24 h. Then increasing doses of doxorubin (Sigma), combined with or without the chelator Trien, were added for 72 h. Medium was removed and cells were incubated with 44 µM resazurin in growth medium for 2h at 37°C. Resorufin was detected using a GloMax 96 Microplate Luminometer (Promega, Southampton, UK)

### Statistical testing

Unless indicated otherwise, for multiple comparisons one-way Anova was used for testing. For two samples, parametric T-tests were used. For Kaplan-Meijer survival curves, Log-Rank test was used where data were studied in quartiles based on expression of the genes of interest. * p<0.05, ** p<0.01, *** p<0.001, **** p<0.0001.

## Supplemental Figure legends

**Figure S1** A) In-Cell western signals intensities of p53 conformational antibodies PAb1620 and PAb240 in the 800 nm channel and of total p53 using antibody FL393 in the 700 nm channel are shown.

**Figure S2** A) Median *SLC31A1* expression in various cancers using the TCGA Pan Cancer dataset. B) Survival of all lung cancer patients in the TCGA Pan Cancer dataset with low or high *SLC31A1* expression. C) Median *MT2A* expression in various cancers using the TCGA Pan Cancer dataset. D) Survival of all lung cancer patients in the TCGA Pan Cancer dataset with low or high *MT2A* expression. E) Proportion of non-smokers, smokers, ex-smokers or unknowns in SCLC or a control group. F) The average age in the SCLC or the control group. SD is indicated. G) Proportion of males and females in SCLC or a control population. H) Copper per amount of tissue measured using ICP-MS in the tumour tissue, adjacent tissue or control samples. The red box indicates the Hamartoma patient. I) Copper per amount of tissue measured using ICP-MS in the serum of SCLC patients or control. The red box indicates the Hamartoma patient. J) Copper per amount of tissue measured using ICP-MS in the smokers and ex-smokers group in the SCLC population. J) Copper per amount of tissue measured using ICP-MS in the smokers and ex-smokers group in the control population. K) Copper per amount of tissue measured using ICP-MS in ex-smokers in the SCLC population or control population.

**Figure S3** A) Immunoprecipitation of p53 using the conformation-specific antibodies PAb1620 and PAb240 in MCF7 cells exposed to CuSO_4_. B) Immunoprecipitation of p53 using the conformation-specific antibodies PAb1620 and PAb240 in HEK293T cells (wt p53) treated with CuSO_4_ in time as indicated. C) Immunoprecipitation of p53 using the conformation-specific antibodies PAb1620 and PAb240 in HEK293T cells (wt p53) treated with CuSO_4_ for 4h, then CuSO_4_ was removed and incubated with copper free medium for in time as indicated for recovery. D) Immunoprecipitation of p53 using the conformation-specific antibodies PAb1620 and PAb240 in HEK293T cells (wt p53) treated with indicated concentrations of CuSO_4_ for 24h in the presence or absence of 100 µM ZnSO_4_ (6h of pre-treatment). E) Immunoprecipitation of p53 using the conformation-specific antibodies PAb1620 and PAb240 of In vitro translated p53. Quantification of 1620 signal over 240 signal is shown at the bottom of this blot. F) Immunoprecipitation of p53 using the conformation-specific antibodies PAb1620 and PAb240 in lysates of A2780 cells exposed to NiSO_4._ G) Cell viability of Beas-2B cells exposed to copper. Error bars indicate SEM. H) Cell viability of A549 cells exposed to copper. Error bars indicate SEM.

**Figure S4** A) Immunoprecipitation of p53 using the conformation-specific antibodies PAb1620 and PAb240 of Hep3B cells transfected with WT p53 and exposed to CuSO_4_ for 8h. p53 was detected by western blot using DO-1 antibody. B) p53 Luciferase reporter assay in Hep3B cells transfected with p53 luciferase and p53 and exposed to copper for 24 h. C) Copper measurements using ICP-MS in Hep3B cells that were incubated in CuSO_4_ for 8h. D) Co-immunoprecipitation of p53 with HA-TAp67α Hep3B cells were transiently transfected with HA-TAp73α in combination with either empty vector (EV), WT, 175H or 273H p53. Hep3B cells were treated with indicated concentrations of CuSO_4_ for 8h and p53 and HA expression were examined using western blot.

**Figure S5** A-B) Proximity ligation assay of p53 with HA-TAp63α (A) or HA-Ago2 (B). Hep3B cells (p53 null) were transiently transfected with HA-TAp63α/HA-Ago2 in combination with WT or 175H p53 and treated with indicated concentrations of CuSO_4_ for 8h.

**Figure S6** Frequencies of the 313 known partially functionally inactivating p53 mutations, by residue, in the TCGA lung cancer (n = 993) and pan-cancer (n = 8741) datasets. Exons 2-11 are labelled numerically below and p53’s domain structure is indicated along the x-axis; TAD = transactivation domain, PRR = proline-rich region, DBD = DNA-binding domain, TET = tetramerization domain, CTD = C-terminal domain.

**Table S1 A:** SCLC patient samples.

| Patient ID | Gender | Smoking status | Age | Tumour | Adjacent | Serum |
| --- | --- | --- | --- | --- | --- | --- |
| 1 | Male | Ex Smoker | 62 | X |  |  |
| 2 | Male | Ex Smoker | 59 | X |  |  |
| 3 | Female | Ex Smoker | 83 | X |  |  |
| 4 | Female | Current Smoker | 63 | X |  |  |
| 5 | Male | Current Smoker | 70 | X |  |  |
| 6 | Female | Current Smoker | 70 | X |  | x |
| 7 | Female | Current Smoker | 47 | X |  | X |
| 8 | Female | Ex Smoker | 74 | X | x |  |
| 9 | Female | Ex Smoker | 59 | X | X |  |
| 10 | Male | Current Smoker | 60 | X | X |  |
| 11 | Female | Ex Smoker | 69 | X | X |  |
| 12 | Female | Current Smoker | 71 | X | X |  |
| 13 | Female | Ex Smoker | 77 | X | X |  |
| 14 | Male | Ex Smoker | 73 | X | X |  |
| 15 | Male | Current Smoker | 74 | X | X | X |
| 16 | Female | Unknown | 77 | X | X | X |
| 17 | Male | Unknown | 75 | X | X | X |
| 18 | Female | Current Smoker | 75 | X | X | X |
| 19 | Female | Current Smoker | 55 | X | X | X |
| 20 | Male | Current Smoker | 69 | X | X | X |
| 21 | Female | Current Smoker | 57 | X | X | X |
| 22 | Male | Ex Smoker | 71 | X | X | X |
| 23 | Female | Ex Smoker | 82 | X | X | X |
| 24 | Male | Ex Smoker | 69 | X | X | X |
| 25 | Female | Ex Smoker | 59 | X | X | X |
| 26 | Male | Ex Smoker | 77 | x | x | X |

**Table S1B:** Benign samples.

| Patient ID | Gender | Smoking status | Age | Diagnosis | Tissue | Serum |
| --- | --- | --- | --- | --- | --- | --- |
| 1 | Female | Unknown | 56 | Aspergillus. No Evidence of Malignancy | X | X |
| 2 | Male | Unknown | 48 | No Evidence of Malignancy | X | X |
| 3 | Female | Unknown | 54 | No Evidence of Malignancy | X | X |
| 4 | Male | Non Smoker | 67 | No information | X | X |
| 5 | Male | Ex Smoker | 64 | No Tumour Identified | X | X |
| 6 | Female | Ex Smoker | 61 | Benign Hamartoma | X | X |
| 7 | Male | Ex Smoker | 67 | Fibrosis and Inflammation | X | X |
| 8 | Female | Ex Smoker | 79 | Mycoplasma Infection | X | X |
| 9 | Male | Ex Smoker | 76 | No Evidence of Malignancy | X | X |
| 10 | Male | Ex Smoker | 71 | No Evidence of Malignancy | X | x |
| 11 | Female | Ex Smoker | 52 | Necrotic Granulomatous Inflammation | x | X |

**Table S2.** Immunoprecipitations.

| **Cells** |  |  |  |
| --- | --- | --- | --- |
| **antibody** | **amount** | **beads** | **size** |
| p53 (DO-1) | 0.2 µg | 30 µl beads | (10cm<sup>2</sup> well) |
| p53 (FL-393) | 1 µg | 30 µl beads | (10cm<sup>2</sup> well) |
| p53 (PAb240) | 0.32 µg | 30 µl beads | (10cm<sup>2</sup> well) |
| p53 (PAb1620) | 0.8 µg | 30 µl beads | (10cm<sup>2</sup> well) |
| HA (16B12) | 2.25 µg | 60 µl beads | (60cm<sup>2</sup> dish) |
| Ago2 (2E12-1C9) | 5 µg | 60 µl beads | (60cm<sup>2</sup> dish) |
| **Explants** |  |  |  |
| **antibody** | **amount** | **beads** | **size** |
| p53 DO-1 | 1 µg | 30 µl beads | (50µg tissue) |
| p53 PAb240 | 1 µg | 30 µl beads | (50µg tissue) |
| p53 PAb1620 | 1 µg | 30 µl beads | (50µg tissue) |

**Table S3.**
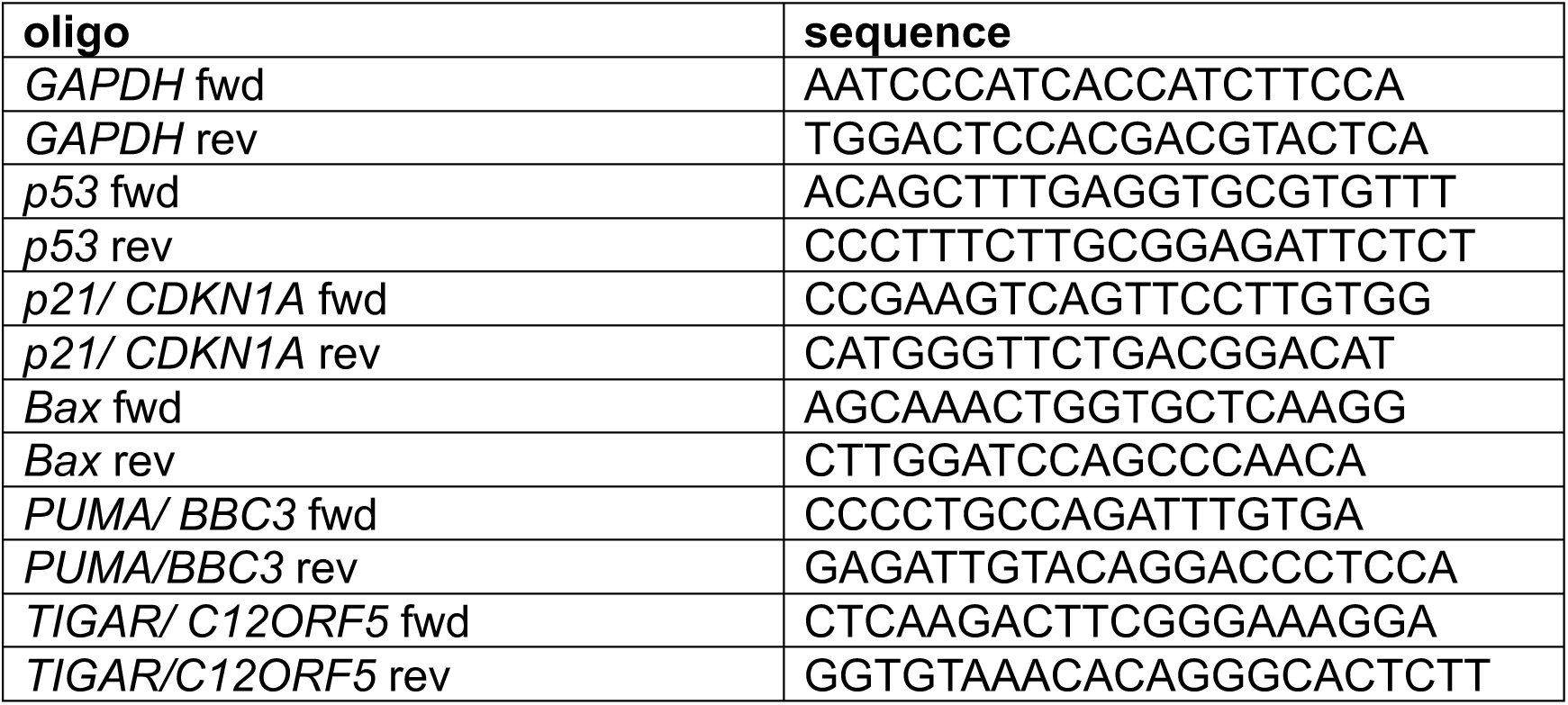
qPCR primers (5’ to 3’)

